# Phycobilisome breakdown effector NblD is required to maintain the cellular amino acid composition during nitrogen starvation

**DOI:** 10.1101/2021.03.18.436103

**Authors:** Vanessa Krauspe, Stefan Timm, Martin Hagemann, Wolfgang R. Hess

**Author notes:** Correspondence: Wolfgang R. Hess.

## Abstract

Small proteins are critically involved in the acclimation response of photosynthetic cyanobacteria to nitrogen starvation. NblD is the 66-amino-acid effector of nitrogen-limitation-induced phycobilisome breakdown, which is believed to replenish the cellular amino acid pools. To address the physiological functions of NblD, the concentrations of amino acids, intermediates of the arginine catabolism pathway and several organic acids were measured during the response to nitrogen starvation in the cyanobacterium *Synechocystis* sp. PCC 6803 wild type and in an *nblD* deletion strain. A characteristic signature of metabolite pool composition was identified, which shows that NblD-mediated phycobilisome degradation is required to maintain the cellular amino acid and organic acid pools during nitrogen starvation. Specific deviations from the wild type suggest wider-reaching effects that also affect such processes as redox homeostasis via glutathione and tetrapyrrole biosynthesis, both of which are linked to the strongly decreased glutamate pool, and transcriptional reprogramming via an enhanced concentration of 2-oxoglutarate, the metabolite co-regulator of the NtcA transcription factor. The essential role played by NblD in metabolic homeostasis is consistent with the widespread occurrence of NblD throughout the cyanobacterial radiation and the previously observed strong positive selection for the *nblD* gene under fluctuating nitrogen supply.

**Importance:** Cyanobacteria play important roles in the global carbon and nitrogen cycles. In their natural environment, these organisms are exposed to fluctuating nutrient conditions. Nitrogen starvation induces a coordinated nitrogen-saving program that includes the breakdown of nitrogen-rich photosynthetic pigments, particularly phycobiliproteins. The small protein NblD was recently identified as an effector of phycobilisome breakdown in cyanobacteria. In this study, we demonstrate that the NblD-mediated degradation of phycobiliproteins is needed to sustain cellular pools of soluble amino acids and other crucial metabolites. The essential role played by NblD in metabolic homeostasis explains why genes encoding this small protein are conserved in almost all members of cyanobacterial radiation.

## Introduction

The recent introduction of the antibiotic retapamulin as an inhibitor of the initiation of translation into ribosome-profiling studies in bacteria has revolutionized the view on the bacterial small proteome suggesting that a high number of previously unexplored small open reading frames (smORFs) exists in bacteria (1, 2). Small proteins, often defined as those containing 50 or fewer amino acids are too small to have own enzymatic activities and this applies also to the group of “larger” small proteins that range from 50 to 100 amino acids. Nevertheless, small proteins have been shown to fulfil multiple important functions, for example as regulatory subunits of multi-protein complexes, as modulators of enzymatic activities of larger proteins or as toxins (for a recent review see Hemm et al. (3)).

A large number of small proteins have been characterized in photosynthetic cyanobacteria due to the extensive work on characterizing the photosynthetic apparatus. Nineteen small proteins with fewer than 50 amino acids are directly involved in photosynthesis. These proteins constitute subunits of photosystem I (encoded by genes *psaM, psaJ* and *psaI* (4)), of photosystem II (genes *psbM, psbT* (*ycf8*), *psbI, psbL, psbJ, psbY, psbX, psb30* (*ycf12*), *psbN, psbF, psbK* (5, 6)), participate in photosynthetic electron transport (Cyt*b*_6_*f* complex; *petL, petN, petM, petG* (7–9)) or in photosynthetic complex I (*ndhP, ndhQ* (10, 11)). Other previously characterized small proteins in cyanobacteria have accessory functions to photosynthesis such as the small CAB like proteins HliA, HliB, HliC and HliD (12–14). The cytochrome *b*_6_*f* complex subunit VIII, encoded by *petN* is with 29 amino acids the shortest annotated photosynthetic protein conserved in cyanobacteria (15). Additional smORFs were computationally predicted in the model cyanobacterium *Synechocystis* sp. PCC 6803 (hereafter designated *Synechocystis* 6803) (16). The 44 amino acids small protein AcnSP was characterized as regulator of the tricarboxylic acid (TCA) cycle enzyme aconitase (17).

Another set of small proteins were found to be involved in the acclimation response of the photosynthetic machinery to nitrogen (N) starvation, including NblA1, NblA2 and NblD, proteins of 62, 60 and 66 residues in *Synechocystis* 6803 (18, 19). Nondiazotrophic cyanobacteria, such as *Synechocystis* 6803, depend on combined nitrogen (N) sources in their environment. These bacteria favor ammonia (NH_3_ or NH_4_^+^) but can also utilize nitrate (NO_3_^-^) as an inorganic N source (20, 21). Under N-limited conditions, cells can start an acclimation program, manifested as chlorosis, and finally enter a dormant state (22–26). During chlorosis, N-rich pigments are degraded, predominantly the giant light-harvesting pigment-protein complexes, the phycobilisomes (22, 23, 27). The excess carbon (C) availability during chlorosis induces the accumulation of C reserves, particularly glycogen (28). The tricarboxylic acid (TCA) cycle intermediate 2-oxoglutarate (2-OG), which is primarily used as a C skeleton for ammonium incorporation by the glutamine synthetase/glutamine 2-OG aminotransferase (GS/GOGAT) cycle (29, 30), has been shown to serve as a cellular signal metabolite to report the relative N/C status (31–35). If ammonia assimilation via the GS/GOGAT cycle is slowed due to N deficiency, 2-OG is no longer converted and accumulates in the cyanobacterial cell. As a result, 2-OG can bind to the signal transduction protein PII (36, 37), thereby releasing PipX from the PipX/PII complex (33, 35). Free PipX can subsequently associate with and activate the global nitrogen transcription regulator NtcA together with 2-OG. In *Synechocystis* 6803, NtcA directly controls 79 different genes after 4 h of N starvation (38). Depending on the location of the respective binding motifs, NtcA can either activate genes, especially those involved in nitrogen uptake and assimilation, or repress genes, as observed for the GS inhibition factors *gifA* and *gifB* (39). Two of the NtcA-activated genes in *Synechocystis* 6803 are *nblA1* and *nblA2* (*ssl0452* and *ssl0453*) (38), which encode small proteins that are critically important in N starvation-induced phycobilisome breakdown (18, 40–42).

Phycobilisome breakdown in N-starved cyanobacteria is a tightly controlled process that is believed to fill cellular amino acid pools, compensating for the limited *de novo* nitrogen assimilation via the GS-GOGAT cycle. NblA1/A2 play essential roles in the coordinated process of phycobilisome degradation during N deprivation because corresponding mutants show a so-called non-bleaching phenotype (*nbl*), i.e., they retain their phycobilisomes under N-limiting conditions (18, 40). NblA proteins function as adaptors for the Clp protease by connecting the phycobiliproteins and the ClpC chaperone (41–44). Phycobilisome degradation can be generally divided into a rapid early phase, in which almost 50% of phycocyanin is degraded within 5 h, followed by a more sedate phase until only approximately 10% of the initial phycocyanin content is attained after 24 h of N deficiency (45). The degradation of the phycobilisome starts at the distal phycocyanin rods and extends to the allophycocyanin core, which is in proximity to the photosystems situated in the thylakoid membrane (44, 46). Previous studies showed that the inability to degrade phycobilisomes in the Δ*nblA1/A2* mutant affected the composition of amino acid pools during N starvation in a specific way in *Synechocystis* 6803 (47). These authors demonstrated that the pool sizes of 12 amino acids were NblA1/A2-dependent, while others were not.

NblD has been recently identified as another small protein determined to be important for phycobilisome turnover because the pigmentation of Δ*nblD* mutants was only marginally reduced under N starvation (19) resembling the non-bleaching phenotype of Δ*nblA* mutants. NblD specifically binds to the phycocyanin beta subunit (CpcB) of phycobilisomes at an early time point of nitrogen depletion (19). However, the precise biochemical function of NbID remains to be determined. In this study, we compared the metabolic composition of *Synechocystis* 6803 wild-type cells (WT) with a mutant deficient for NblD (Δ*nblD*) over a time course of N starvation. A characteristic signature in the composition of these metabolite pools was observed that supports the essential role played by NblD in the maintenance of cellular amino acid homeostasis during N starvation. Other deviations from the WT suggest wider-reaching effects, also affecting transcriptional reprogramming via an unusually high concentration of 2-OG, the metabolite co-regulator of the NtcA transcription factor.

## Results

### Global responses of an *nblD* deletion mutant to nitrogen starvation

In the present study, the amino acid pools and amounts of other important metabolites were compared in the *Synechocystis* 6803 wild type (WT) to an *nblD* deletion mutant (Δ*nblD*). In the absence of a nitrogen source, cells of the Δ*nblD* strain are bleaching significantly less than the WT (19). Accordingly, we hypothesized that the soluble amino acid pools of Δ*nblD* cells may be exhausted more quickly than those of WT cells. This assumption was tested by metabolomics based on liquid chromatography mass spectrometry (LC-MS/MS) of three biologically independent samples for each WT and Δ*nblD*, taken before (+N condition), and 3 h or 24 h after the transfer into N-free BG11 medium. In total, 28 compounds, including almost all amino acids (Asn and Cys were below the detection limit), intermediates of the arginine catabolism pathway, and several organic acids, were quantified (see Table S1 for the LC-MS/MS raw data).

Cells obtained from the +N condition and 3 h after nitrogen removal showed no difference in pigmentation. After 24 h of N starvation, the WT samples showed pronounced bleaching, while the Δ*nblD* mutant bleached much less (Fig. 1A and B). Comparing the spectra with previously published spectra suggests that the decrease in the 630 nm absorption maximum of phycobilisomes in Δ*nblD* under N starvation is greater than in the Δ*nblA1/nblA2* double mutant (18, 42), although clearly not as much as in the WT under the same condition. The degradation of phycobilisomes contributed to transiently higher levels of total soluble amino acids in *Synechocystis* 6803 WT during N starvation (Fig. 2A, Table S2). The level of free amino acids increased from approximately 261 ng OD_750_^-1^ ml^-1^ (± 32.4) under nitrogen-replete conditions (+N) by 65% on average after 3 h and approximately 27% (in relation to the starting value) after 24 h of N limitation (-N). In contrast, the Δ*nblD* mutant contained 335 ng OD_750_ ^-1^ ml^-1^ (± 43.3) of free amino acids in the +N condition and remained at a similar level after 3 h. After 24 h after nitrogen removal, the amino acid content decreased dramatically by 70% to 75% in relation to the initial content in Δ*nblD* (Fig. 2A). Compared to the amino acid pool, the total amounts of other quantified metabolites, primarily organic acids, did not differ significantly between cells of the WT and Δ*nblD* (Fig. 2B), at least in the +N condition and after 24 h. The higher total amount after 3 h is mostly due to increased 2-OG and citrate concentrations in the mutant (see below).

**Figure 1.**
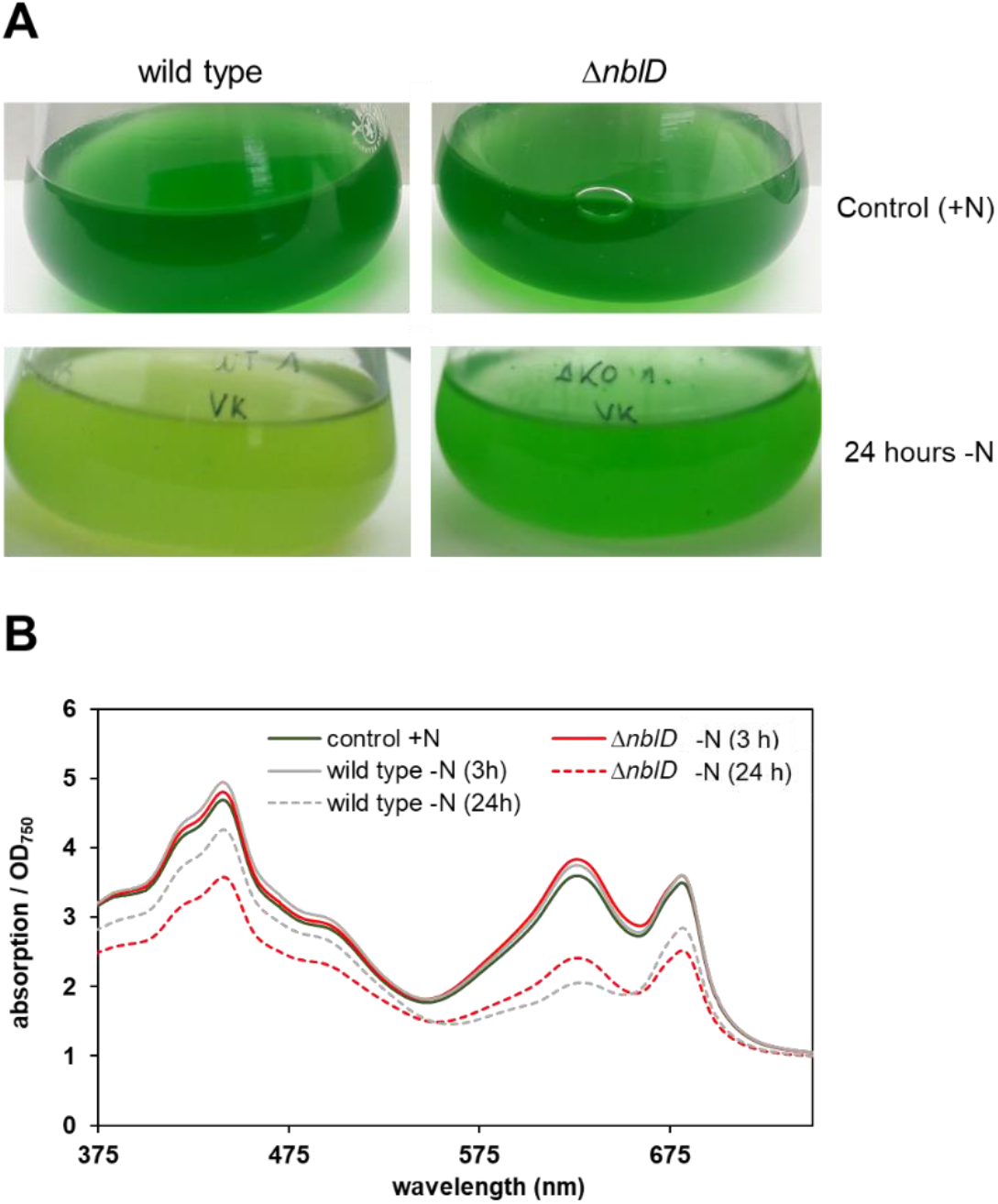
Response of the *Synechocystis* 6803 WT and Δ*nblD* mutant during the response to N starvation. **A**. Phenotype under nitrogen-replete (+N) conditions and after 24 h of nitrogen depletion (-N). **B**. Spectrometric measurements of cultures normalized to OD_750_.

**Figure 2.**
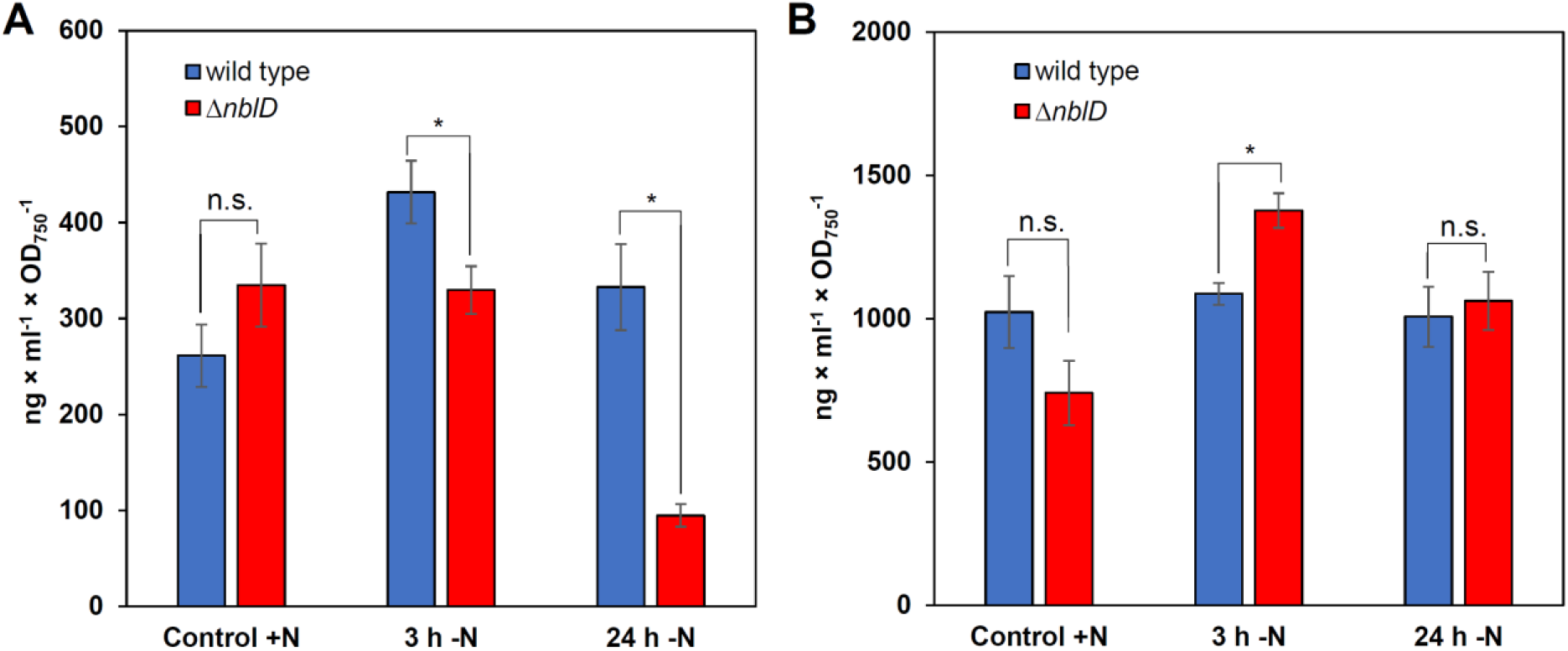
A. Total soluble amino acid content and **B**. total soluble metabolite content (non-proteinogenic amino acids and important intermediates) of wild-type (blue) and Δ*nblD* (red) cultures in nitrogen-replete (+N) and nitrogen-depleted BG11 (-N) after 3 h and 24 h. For each time point and strain, triplicates were measured, and significance (p < 0.05) was calculated by t-test; n.s. = not significant. The raw data are given in Table S2.

### Changes in the composition of amino acid pools differ between WT and Δ*nblD*

Despite the overall trend of a decreasing total amino acid pool in mutant Δ*nblD* compared to WT, the direction of changes in the amounts of individual amino acids were not unidirectional (Fig. 3). According to their differential changes between WT and Δ*nblD* during the N starvation period, these changes could be categorized into four groups. Group 1 comprised amino acids showing no major differences between WT and Δ*nblD;* group 2 included those amino acids that appeared different already at the +N condition, while group 3 exhibited differences between WT and Δ*nblD* early, after 3 h, or group 4 later, after 24 h of N starvation (Fig. 3).

**Figure 3.**
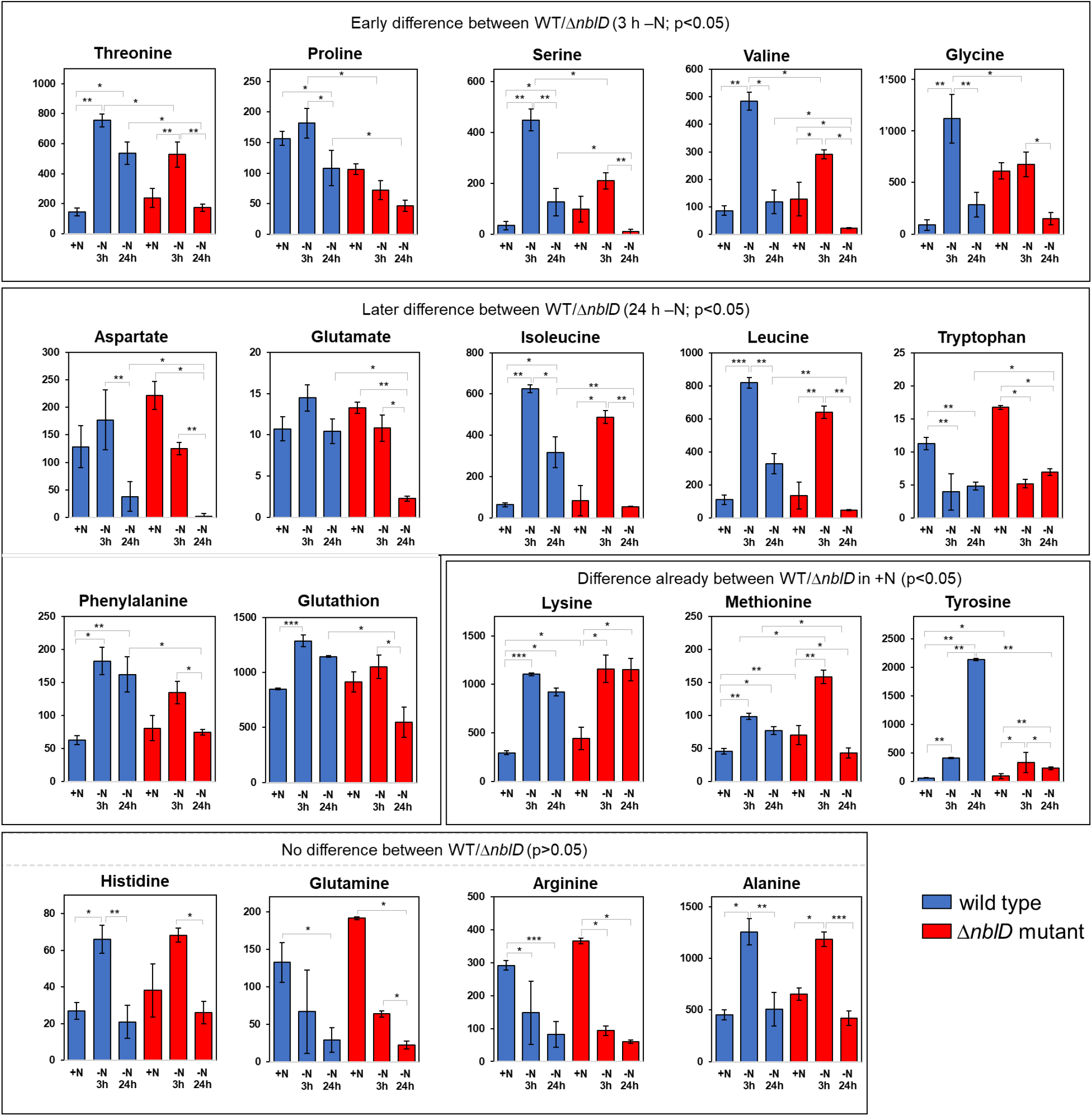
Development of amino acid content (in mM for Glu and µM for all other) in WT (blue) and Δ*nblD* mutant (red) in nitrogen replete (+N) and nitrogen deplete (-N) medium after 3 h and 24 h. Amino acids are grouped together according to the observed differences between wild type and mutant as follows: Difference already observed in +N conditions; no difference between strains; early difference (after 3 h); later differences (24 h). Significance was calculated with a two-sample t-test with unequal variance (Welch’s t-test; p < 0.05 = *; p < 0.01 = ** and p < 0.001 = ***) for the development in each strain and between the strains at corresponding time points (see Table S4 for details).

Amino acids of group 1 showed similar trends in both WT and Δ*nblD* during nitrogen limitation, as was observed for Ala, Arg, Gln and His. The strongest deviation in the response to N starvation between the two strains was observed for Glu. The level of Glu did not decrease during the 24 h N starvation period in WT but decreased in mutant Δ*nblD* to approximately 17% of the initial value after 24 h of N starvation (Fig. 3). Glu is by far the most abundant soluble amino acid and serves as an amino group donor in the majority of transaminase reactions. Hence, the depletion of the Glu pool should have notable effects on the entire metabolism. Indeed, the lack of Glu extended into the pool of reduced glutathione (GSH), which is the tripeptide γ-L-glutamyl-L-cysteinylglycine; the initial increase observed in the WT was lower in cells of the mutant Δ*nblD* and led after 24 h into N limitation to a significantly reduced amount (Fig. 3). GSH plays an important role as a reducing agent in all living cells and was found to be essential in the acclimation of *Synechocystis* 6803 to environmental and redox perturbations (48); hence, its depletion probably should have an impact on redox homeostasis in the mutant Δ*nblD*. The amino acids of group 3, such as Gly, Pro, Ser, Thr and Val, exhibited transiently increased levels after 3 h in WT, which was not the case or was less pronounced in the mutant (Fig. 3). The branched chain amino acids Ile, Leu and Val share the same biosynthetic pathway, whereas Gly and Ser are closely linked to photorespiratory metabolism and can be interconverted by serine hydroxymethyltransferase (49). Phe and Thr exhibited increased levels in WT cells after 24 h of N starvation, while they returned to initial levels in Δ*nblD*, accumulating at values of less than 50% of the WT. Finally, the aromatic amino acid Tyr from group 2 accumulated nearly 100-fold more in WT cells after 24 h of N starvation, whereas it remained at low levels in Δ*nblD* (Fig. 3). This finding is consistent with a previous analysis of *Synechocystis* 6803 WT after 24 h N starvation (47). Tyr represents approximately 5% of the amino acids stored in the phycobilisome (Fig. 4). The strong accumulation of Tyr in WT but not in Δ*nblD* suggests that it originated directly from phycobilisome degradation, assuming that in contrast to other amino acids, Tyr is not readily incorporated into the metabolism of N-starved cells.

**Figure 4.**
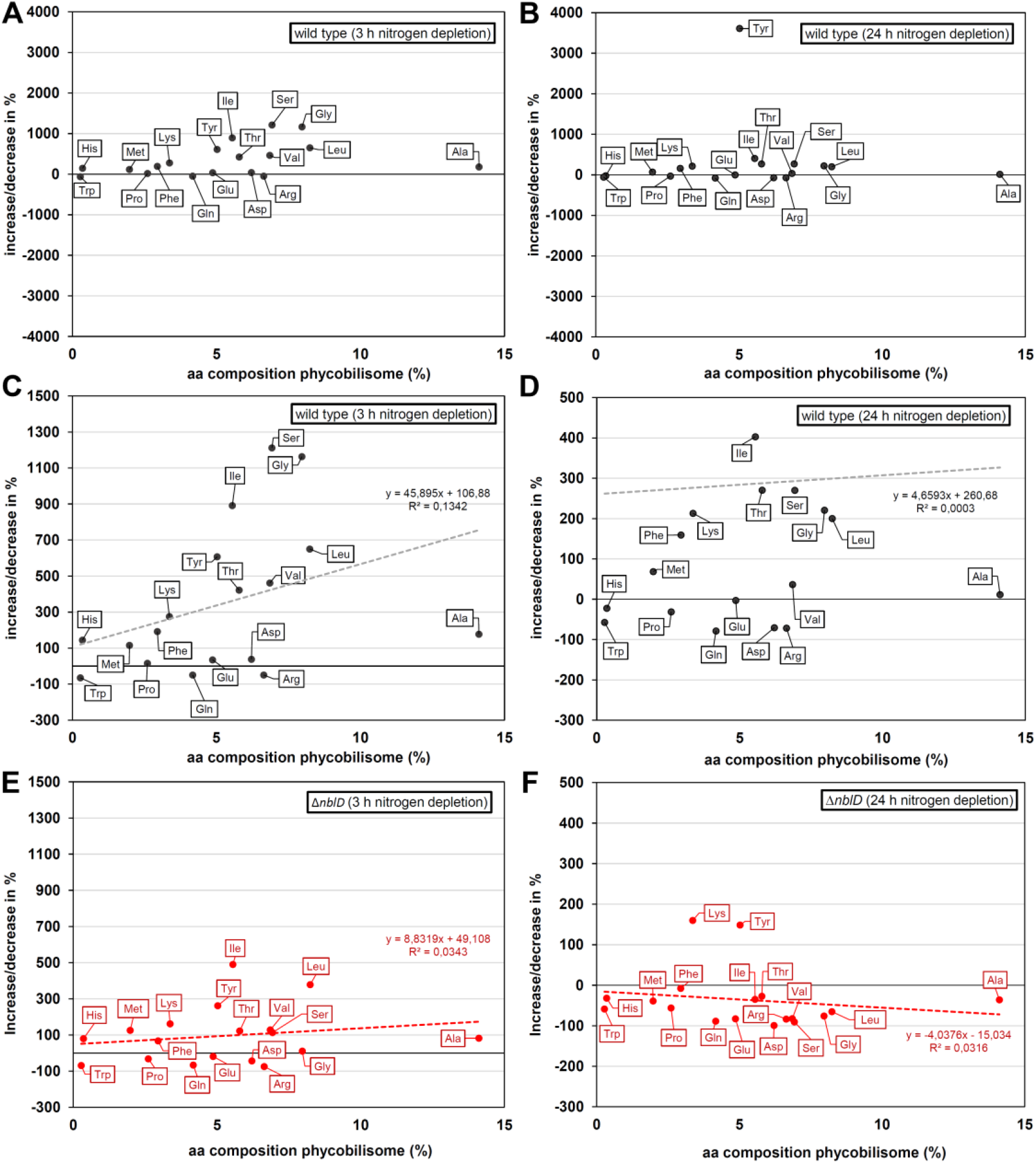
Phycobilisome individual amino acid (aa) composition (in %) plotted against their increase or decrease observed in relation to the initial values measured in nitrogen sufficient conditions. **A, C**. WT after 3 h of N limitation; **B, D**. WT after 24 h of N limitation. **E, F**. Data for Δ*nblD* mutant after 3 and 24 h. Phycobilisome composition was calculated based on the structure description by Arteni et al. (55), assuming a common complex with 6 rods each with 3 hexameric phycocyanin discs and a tricylindrical allophycocyanin core. A linear trend line shows very low correlation between the variables, and a poor R-squared value indicates high variation of the data in relation to this line.

### Changes in other N-containing metabolites and organic acids

In addition to proteinogenic amino acids, many additional metabolites could be quantified. For example, ornithine, a key intermediate in the arginine catabolism pathway, was clearly depleted in Δ*nblD* upon N starvation compared to WT (Fig. 5). This points to the arginine catabolism pathway as another mechanism for stabilization of the Glu pool. In this pathway, the arginine-guanidine-removing enzyme (AgrE) together with proline oxidase (PutA) can produce Glu from the Arg molecules released from phycocyanin via ornithine (50–52). Therefore, the diminished ornithine pool adds another important element to the changed nitrogen homeostasis in cells of the Δ*nblD* strain. This assumption was also supported by changes in the 2-OG pool, which was lower in WT than in the mutant under all conditions, indicating a relatively more N-starved state of Δ*nblD* cells. Moreover, we observed that the absence of NblD also affected the pool of 3-phosphglyceric acid (3PGA), the stable CO_2_ fixation product of RubisCO, which was decreased under all conditions. Excess 3PGA can be used to produce glycogen, which is highly accumulating in N-starved cells during chlorosis (53). Otherwise, some 3PGA can be diverted via phosphoglycerate mutase into the direction of lower glycolysis and the TCA cycle, which is primarily regulated by PirC (54). In contrast to the decreased flow into the oxidative branch, the reductive branch of the TCA cycle seemed to be less affected by the absence of NblD, since the pool of malate was similar in cells of the mutant and the WT (Fig. 5). However, succinate was clearly lowered in Δ*nblD* after 3 h under N limitation, probably due to altered consumption of succinate by the respiratory chain. In the mutant cells, the succinate pool stabilized at a similar quantity as was observed in WT cells under prolonged N depletion. Taken together, the metabolic changes observed in this study enabled us to conclude that the absence of NblD had a broad impact on primary metabolism.

**Figure 5.**
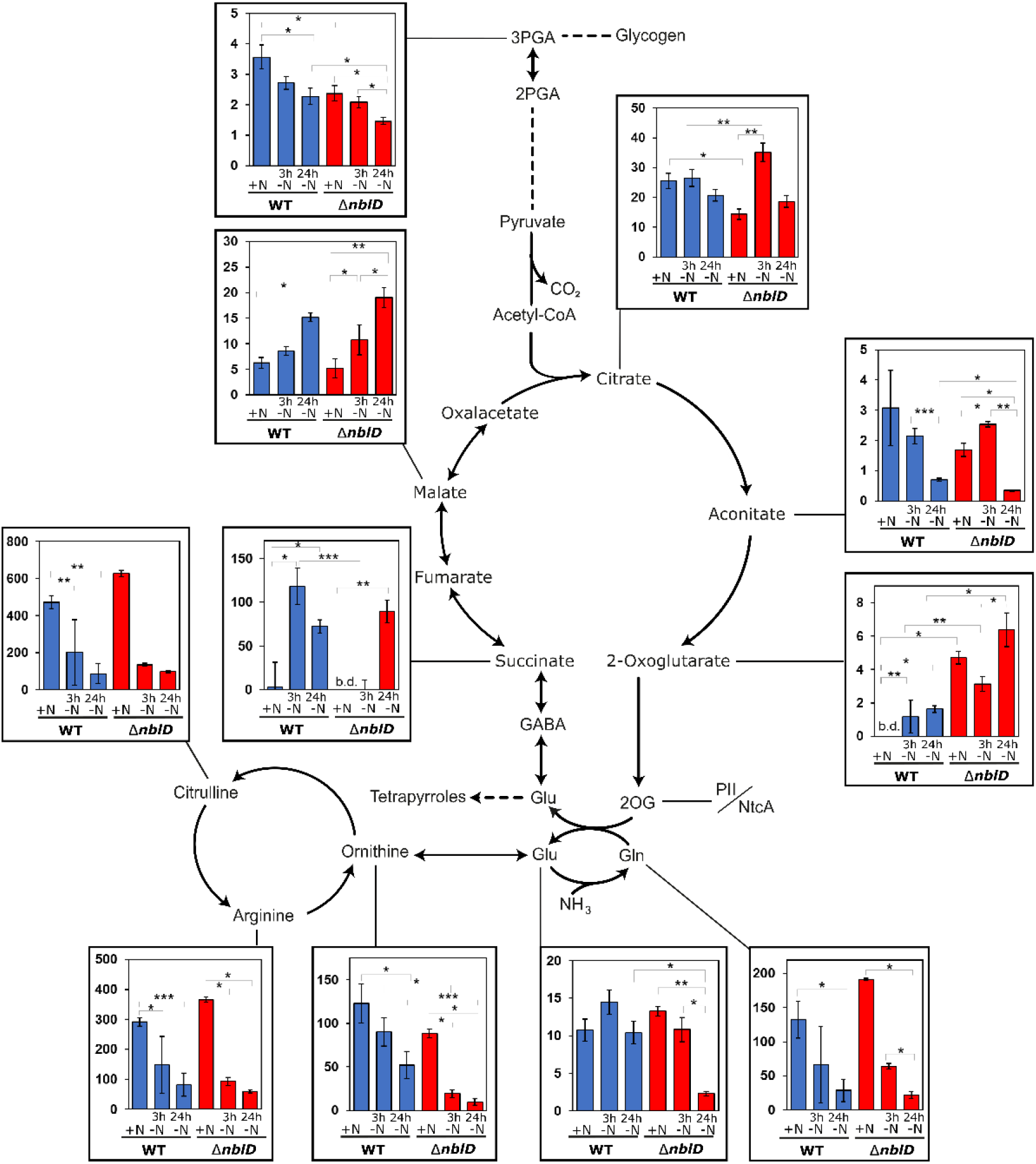
Comparison of metabolite content (in mM for Glu, citrate, 3PGA, malate and 2-OG, in µM for all other) in WT (blue) and Δ*nblD* deletion mutant (red) in nitrogen replete (+N) and nitrogen deplete (-N) medium after 3 h and 24 h shown alongside their biosynthetic pathways. Succinate was below detection limit (b.d.) in the Δ*nblD* sample under nitrogen-replete conditions. Significance was calculated with the two-sample t-test with unequal variance (Welch’s t-test; p < 0.05 = *; p < 0.01 = ** and p < 0.001 = ***) for the development in each strain and between the strains at corresponding time points (details in Table S4).

## Discussion

### Comparison of the changes in Δ*nblD* with the previously characterized Δ*nblA1/A2* mutant

NblD is involved in the disassembly of phycobilisomes by targeting CpcB during acclimation to N starvation (19). In contrast to the WT, the Δ*nblD* mutant maintains the majority of its pigments under this condition resembling the non-bleaching phenotype of the previously reported Δ*nblA* strains and suggesting that phycobilisomes are retained in these mutants to a large extent during the considered time frame (see Fig. 1). In case of the Δ*nblA1/A2* mutant, amino acid pools were analyzed after 24 h in the presence or absence of nitrogen. The authors categorized the changes in the amino acid pool sizes into NblA1/A2-dependent and -independent responses. Most amino acids (Gln, Glu, Gly, Ile, Leu, Met, Phe, Pro, Ser, Thr, Tyr, Val and also GSH) fell into the NblA1/A2-dependent category. In the present study, we analyzed the effects of the absence of NblD on the amino acid and organic acid pools in the Δ*nblD* deletion mutant after 3 h and 24 h of N starvation. The observed differences between Δ*nblD* and WT in the amino acid pools were similar to those described for the Δ*nblA1/A2* mutant (47) but were not identical, despite the fact that both types of proteins play a role in phycobilisome degradation. Five amino acids (Ile, Leu, Ser, Thr, Val) had after 24 h of N starvation higher levels in WT but lower levels in the Δ*nblA1/A2* mutant compared to the control in N-replete media. While we observed the same changes for these 5 amino acids after 24 h N starvation, we noticed that their pools were peaking even higher in WT in the sample taken after 3 h. In the study by Kiyota et al. (47), the WT had been analyzed at even higher temporal resolution (after 1, 3, 6, 12 and 24 h of N starvation) and they reported that these 5 amino acids transiently overshot at the 3 h value, hence our data are consistent. However, we found a transient accumulation of Ile, Leu, Ser, Thr, and Val in cells of the Δ*nblD* mutant like in the WT in the 3 h sample. Hence, our data allow to differentiate between an early NblD-independent response for these 5 amino acids and a later response that was NblD-dependent. This finding suggests that the early, very pronounced response was independent from amino acid release from the phycobilisomes, whereas they were mobilized at the later time point, which both the Δ*nblD* and Δ*nblA1/A2* mutants failed to do.

The levels of four other amino acids (Pro, Gly, Glu, Tyr) and GSH behaved very similar in the Δ*nblA1/A2* and Δ*nblD* mutants. However, for three amino acids (Gln, Met and Phe) we observed differences between the different non-bleaching mutants. Gln levels steadily declined in our study, both in WT and Δ*nblD* over the time of the experiment; hence, it appeared as an NblD-independent response (Fig. 3). For Met and Phe, similar or higher concentrations had been reported in the WT but lower in Δ*nblA1/A2* upon N starvation, with Met even falling below the detection limit after 24 h (47). While we observed the same effect for the WT, in contrast the Δ*nblD* mutant exhibited transiently higher levels of Met and Phe after 3 h, and about the same levels after 24 h as at the beginning.

In the work by Kiyota et al. (47) some amino acids were identified as NblA1/A2-independent (Ala, Asn, Asp, Lys and Trp). The changes in the pools sizes of these amino acids were also NblD-independent in our work. We can add Arg and His to this category, which were below detection limit in the work on NblA, while we could not measure Asn. Moreover, the decline in Gln levels appeared as an NblD-independent response, different from the findings on NblA.

Collectively, the overall responses of Δ*nblA1/A2* and Δ*nblD* mutants were similar but some deviations appeared. For example, our investigation of the 3 h time point allowed to conclude that some amino acid changes are independent from amino acid releases via the phycobilisome degradation. Consequently, though NblA1/NblA2-and NblD-assisted phycobilisome degradation represents the most obvious protein degradation process due to visible pigment changes, faster changes in the proteome likely contribute to the dynamics of amino acid changes especially during the early phase of acclimation to N starvation.

### Phycobilisome proteins and amino acid pools during nitrogen starvation

Degradation of phycocyanin rods and their proteolytic recycling are assumed to contribute extensively to the soluble amino acid pool in the cell, making those amino acids available to be incorporated into other proteins to acclimate towards nutrient limitation. From the phycobilisome core, up to six rod antennas radiate, each of which is built by three hexameric discs connected with linker polypeptides CpcG and CpcC (55). One hexameric phycocyanin disc contains six subunits of each CpcA and CpcB.

Therefore, the most abundant phycobilisome proteins are CpcA and CpcB, which are rich in Ala, Gly, Leu, Val and Tyr. Assuming full phycobilisome turnover, including also all linker proteins and allophycocyanin core, it would provide mostly Ala (∼14%), Leu, Gly (both ∼8%), Ser, Val, Arg, and Asp (contributing 6-7% each to phycobilisome composition). Accordingly, the pools of these amino acids were expected to increase early during N-starvation-induced phycobilisome degradation in the WT but not in Δ*nblD*. Such an increase was clearly observed for Tyr, whereas the changes in the amounts of the other amino acids differed less pronounced between WT and Δ*nblD* cells. The amounts of Gly, Leu, Ser, and Val were always higher in the WT than in the mutant under -N conditions. However, the amount of Ala, the most frequent amino acid in the phycobilisome showed a similar pool change in WT and mutant cells, suggesting its increase resulted from synthesis instead. Ala biosynthesis is derived from the transamination of pyruvate and Glu by alanine dehydrogenase (*sll1682*) or by serine-pyruvate aminotransferase using pyruvate and Ser. Therefore, Ala is directly linked to the metabolic pathways of Ser, Gly and Thr, as well as to primary C metabolism, via pyruvate. Indeed, in *Synechococcus elongatus* PCC 7942, alanine dehydrogenase was found to be required for correct phycobilisome degradation during N starvation, resulting in lower *nblA* transcription if the gene for alanine dehydrogenase was deleted (56).

As mentioned above, besides Tyr the largest difference between WT and mutant Δ*nblD* was observed for the pool of Glu after 24 h of N starvation. Glu is can be regarded as the central amino acid from several perspectives. It is certainly the most relevant amino group donor for transaminations, consistent with its abundance, which is far higher than any other amino acid. In addition, Glu serves as the precursor of tetrapyrrole synthesis, leading to cytochromes and especially chlorophylls in cyanobacteria, green algae and plants upon its activation by binding to its cognate tRNA^glu^ (57–59). This C5 pathway is the sole biosynthetic pathway for tetrapyrrole biogenesis in *Synechocystis* 6803 (60, 61). Glu has an approximate 5% share in the composition of the phycobilisome proteins (Fig. 4). Therefore, it is likely that amino groups of abundant amino acids that become available upon phycobilisome degradation are utilized predominantly to stabilize the Glu pool via transaminase reactions. Initially, stabilization of the Glu pool may occur through glutamate synthase from Gln and 2-OG, which is also consistent with the relatively lower amounts of 2-OG in WT compared to Δ*nblD* cells (Fig. 5), then via arginine catabolism from Arg. A corresponding reduction in Gln and Arg pools is seen at 3 h and at 24 h into N starvation in both WT and Δ*nblD*. At the later time point, additionally Asp, Gly, Leu, possibly Ala and Val, might be a source of amino groups as these amino acids are (i) relatively abundant, (ii) relatively frequent in the phycobilisome and (iii) their pools initially increased in the WT after shift to N starvation, followed by a particularly sharp reduction from 3 h to 24 h. In addition, the development of pool sizes for these amino acids showed marked differences between the WT and Δ*nblD*.

Our results support the hypothesis that amino acid pools are regulated to a large extent independently of phycobilisome degradation by NblA1/A2 and NblD, because the cells maintain distinct amino acid pool homeostasis involving transaminase reactions, primarily using the amino groups from the released phycobilisome constituents. Furthermore, such amino acids as Ala, Asp and Lys might originate from the breakdown of several other proteins (47). Hence, even though NblA1/NblA2-and NblD-assisted phycobilisome degradation represents the most obvious protein degradation process due to visible pigment changes, additional changes in the proteome likely contribute to the recycling of amino acids during acclimation to N starvation.

### Effects on the organic acids pools

The striking differences between WT and Δ*nblD* observed for Glu at 24 h of N starvation may extend also further, into the amounts of certain organic acids, especially succinate. At 3 h -N, succinate levels were very low, whereas Glu levels were high; at 24 -N, succinate levels were relatively high, whereas Glu levels were very low. There are four possibilities for the relatively high succinate levels after 24 h of N starvation:

i. it may appear from an enhanced flux into the reductive branch of the TCA cycle working in anabolic direction in both strains, because the malate content also increased over time;
ii. an enhanced closure of the “open” cyanobacterial TCA cycle due to the shunt, which converts 2-OG into succinate via 2-oxoglutarate decarboxylase and succinic semialdehyde dehydrogenase reactions (62) would provide higher succinate amounts;
iii. Succinate could also originate from the gamma-aminobutyric acid shunt closing the “open” TCA cycle in which glutamate is converted into gamma-aminobutyric acid and then via succinic semialdehyde to succinate (63);
iv. The succinate consumption by respiration via succinate dehydrogenase could have become reduced.

The flux through the alternative TCA shunts has been regarded to be rather low under light conditions (62, 63) as applied here, therefore, the possibility (ii) appears rather unlikely. Possibility (iv) is unlikely as respiration does not decrease but increases upon N limitation (24). Consequently, the most likely scenario is that the succinate accumulation after 3 h N starvation in WT cells resulted from increased flow through the arginine catabolism pathway from phycobiliproteins (variant iii) via Glu, while the succinate accumulation after 24 h in the WT and especially pronounced in Δ*nblD* originates from the reductive branch of the TCA cycle (i). This scenario is supported by the N starvation-induced expression (53, 64) of *slr1022* encoding an enzyme that functions also as γ-aminobutyrate aminotransferase (63), a key enzyme in the gamma-aminobutyric acid shunt pathway interconverting Glu to succinate.

### Connecting metabolic changes and the regulation of gene expression

Several of the metabolites showing marked changes between WT and mutant Δ*nblD* possess signaling functions connecting metabolism and the regulation of gene expression. For example, Glu is used as a signaling molecule in green algae, mosses and higher plants sensed by the PII protein (65). Moreover, in several Gram-negative bacteria, Gln serves as a signaling molecule for N availability (reviewed by Forchhammer, 2013 (66)). More recently, it was observed that Gln also exhibits a signaling function in *Synechocystis* 6803 and other cyanobacteria via interaction with the GlnA riboswitch governing the expression of *gifB* mRNA (67).

In contrast to Gram-negative bacteria, cyanobacteria use the TCA cycle intermediate 2-OG to sense the C/N status. 2-OG is the regulatory co-factor of NtcA and of PII, where the 2-OG accumulation transduces the low N signal leading to the activation of NtcA (36, 37). Conversely, under high N levels 2-OG is converted via the GS/GOGAT cycle to Glu, thereby the low 2-OG level diminishes the activity of NtcA. In addition, the decrease in 2-OG is characteristic for C limitation and activates the transcription factor NdhR to upregulate genes involved in the cyanobacterial inorganic C-concentrating mechanism (68, 69).

Hence, the increase in 2-OG in the WT observed during N starvation should represent a pronounced signal for activation of NtcA, and indeed, the transcriptional units within the NtcA regulon become strongly differentially regulated during this condition (19). Notably, there is an NtcA motif in the promotor region of the *nblD* gene itself (16), consistent with its role in N-limited conditions. However, compared to WT, we observed that 2-OG increased even more in Δ*nblD* and at every sampling point, including the +N condition (Fig. 5). Because of the function of 2-OG as the main cellular indicator metabolite to report C/N homeostasis (31–35), increased 2-OG levels therefore clearly indicated a reduced ammonia assimilating capacity of mutant Δ*nblD* already under N replete conditions. The likely reduced use of 2-OG in the GS/GOGAT cycle was supported by the observation that the levels of citrate and aconitate, precursors of 2-OG in the oxidative branch in the open cyanobacterial TCA cycle, were lowered in Δ*nblD* (Fig. 5). If the flux into the GS/GOGAT cycle would be similar, this decrease should extend into the 2-OG pool; however, its increase indicates reduced 2-OG utilization in mutant cells, whereas 2-OG was extensively used in WT to replenish the Glu pool.

It could be expected that the higher 2-OG level in Δ*nblD* than in the WT cells would contribute to the stimulation of the NtcA regulon, which was observed in Δ*nblD* during the previous transcriptome analysis. Indeed, 3 h after the induction of N starvation, the mRNA levels of NtcA-regulated *nblA1A2* genes were more strongly induced compared to the WT (log_2_ fold changes of +4.28 and +4.09 in Δ*nblD* and +3.71 and +3.47 in the WT), while *gifA* and *gifB* were more strongly reduced (for both genes, log_2_ fold change of −4.7 in Δ*nblD* and −2.4 in the WT control) (19). By comparison, 2-OG concentrations were around 3.14 mM and 1.16 mM in Δ*nblD* and WT, respectively (Table S3). Hence, a higher 2-OG concentration corresponded roughly to higher mRNA levels 3 h after the induction of N starvation. However, in the nitrogen-replete condition we measured an ∼100 fold higher level of 2-OG in Δ*nblD* compared to WT but the *nblA1A2* mRNA levels were only marginally higher. This result strongly suggests that the high accumulation of 2-OG alone is not sufficient to induce the complete N starvation response and an additional signal can be postulated. Furthermore, the high accumulation of 2-OG in Δ*nblD* already in the +N condition can be taken as evidence that NblD, in addition to its prominent function during N starvation, also plays some role under N-replete conditions. This function of NblD could be related to the remodeling of phycobilisome composition, size and number per thylakoid membrane or to the turnover of phycobilisomes occurring under N-replete conditions.

## Materials and Methods

### Cultivation conditions and sampling for metabolite analysis

*Synechocystis* 6803 PCC-M was used as the WT and background for the construction of mutants. *NblD* knockout mutants were generated by homologous replacement of the *nblD* coding sequence with a kanamycin resistance cassette (*nptII*) (19).

Strains were maintained in copper-free BG11 supplemented with 20 mM TES PUFFERAN^®^, pH 7.5, under continuous white light of 50 µmol photons m^-2^ s^-1^ at 30°C.

Wild-type and Δ*nblD* cultures were grown to OD ∼0.8 in triplicate without antibiotics before cells were pelleted by centrifugation (3,210 × *g*, 8 minutes, room temperature), washed three times in BG11 without N and resuspended in BG11 lacking a N source. After three and 24 h of incubation in N-limited conditions, the OD was recorded, and 2 ml of each culture was collected by centrifugation (9,000 × *g*, 1 minute, room temperature) and immersed in liquid nitrogen. As controls, samples were also obtained before N removal (+N). For extraction, 630 µl methanol containing carnitine (internal standard, 1 µg per extraction) was added, mixed for one minute and incubated for 5 minutes in a sonic water bath. After another 15 minutes of shaking at room temperature, 400 µl of chloroform was added, and the sample was incubated at 37°C for 5 minutes. Next, 800 µl ROTISOLV® LC-MS grade H_2_O (Roth) was added, and the sample was mixed thoroughly. Precipitation was enabled by storage at −20°C for at least 2 h, and phase separation was subsequently achieved by centrifugation for 5 minutes at room temperature (16,800 × *g*). The upper polar phase was collected and dried in a speed vac for 30 minutes followed by lyophilization overnight. A mock sample without any culture was subjected to the same treatments as the other samples.

### Metabolite analysis

Absolute metabolite contents were quantified on a high-performance liquid chromatograph mass spectrometer LCMS-8050 system (Shimadzu) as described previously (70). Dried samples were dissolved in 200 µl LCMS grade water and filtered through 0.2-mm filters (Omnifix-F, Braun, Germany), and 4 µl of the cleared supernatant was separated on a pentafluorophenylpropyl column (Supelco Discovery HS FS, 3 mm, 150 3 2.1 mm).

The compounds were identified and quantified using the multiple reaction monitoring (MRM) values given in the LC-MS/MS method package and the LabSolutions software package (Shimadzu). Authentic standard substances (Merck) at varying concentrations were used for calibration, and peak areas were normalized to signals of the internal standard (carnitine). The data were further normalized to the OD_750_ measured for each sample and corrected by signals obtained from the mock sample. The raw data and estimations as absolute values in nanogram × ml^-1^ × OD_750_ ^-1^ are given in Table S1, statistical evaluations in Table S4. These data were converted to molar concentrations of metabolites per total cell volume assuming an average cell diameter of 3 µm and 10^7^ cells per milliliter at OD_750_ = 1 (Table S3).

## Author Contributions

WRH designed the project and secured funding. VK constructed and cultivated strains, MH and ST measured metabolite concentrations. VK, MH and WRH wrote the manuscript with input from all authors.

## Supplementary Tables

**Table S1**. LC-MS/MS data for 18 amino acids (Cys and Asn are missing because they were below detection limit), several organic acids and GSH. Retention time (r) was individually monitored with 2 standards (line 4 and 5) and the average peak area for 1 ng (line 7), which were used to determine the ng per sample. The carnitine standard (columns E, F, G, H) was used to generate the Factor IS (internal standard), which is implemented in the normalization of the peak area for each sample.

**Table S2**. Total amount of all amino acids and the other soluble metabolites investigated in this study. Average, standard deviation and significance calculated by two-sample t-test (Welch’s t-test) are visualized in figure 2.

**Table S3**. Conversion in molar concentrations of metabolites in µM (Glu in mM) per total cell volume assuming an average cell diameter of 3 µm and 10^7^ cells per milliliter at OD_750_ = 1.

**Table S4**. Welch’s t-test (two-sided, inequal variation) for evaluating significant differences in the amino acids and organic acids between the strains and the development in the strains. Significant differences are marked in red (p < 0.05), boxed (p < 0.01) and bold italics (p < 0.001).

